# On the replicability of diffusion weighted MRI-based brain-behavior models

**DOI:** 10.1101/2024.07.08.602202

**Authors:** Raviteja Kotikalapudi, Balint Kincses, Giuseppe Gallitto, Robert Englert, Kevin Hoffschlag, Jialin Li, Ulrike Bingel, Tamas Spisak

## Abstract

Establishing replicable inter-individual brain-wide associations is key to advancing our understanding of the crucial links between brain structure, function, and behavior, as well as applying this knowledge in clinical contexts. While the replicability and sample size requirements for anatomical and functional MRI-based brain-behavior associations have been extensively discussed recently, systematic replicability assessments are still lacking for diffusion-weighted imaging (DWI), despite it being the dominant non-invasive method to investigate white matter microstructure and structural connectivity. We report results of a comprehensive evaluation of the replicability of various DWI-based multivariate brain-behavior models. This evaluation is based on large-scale data from the Human Connectome Project, including five different DWI-based brain features (from fractional anisotropy to structural connectivity) and 58 different behavioral phenotypes. Our findings show an overall moderate replicability, with 24-31% of phenotypes replicable with sample sizes of fewer than 500. As DWI yields trait-like brain features, we restricted the analysis to trait-like phenotypes, such as cognitive and motor skills, and found much more promising replicability estimates, with 67-75% of these phenotypes replicable with n<500. Contrasting our empirical results to analytical replicability estimates substantiated that the replicability of DWI-based models is primarily a function of the true, unbiased effect size. Our work highlights the potential of DWI to produce replicable brain-behavior associations. However, it shows that achieving replicability with small-to-moderate samples requires stable, reliable and neurobiologically relevant target phenotypes. Our work highlights the potential of DWI to produce replicable brain-behavior associations, but only for stable, reliable and neurobiologically relevant target phenotypes.

**HIGHLIGHTS:**

1. **Moderate replicability in DWI-based models:** Overall replicability of DWI-based brain-behavior associations ranges from 24-31% with sample sizes under 500.
2. **Improved replicability for trait-like phenotypes:** Trait-like phenotypes e.g., cognitive and motor skills exhibit higher replicability estimates of 67-75%, compared to state-like phenotypes such as emotion.
3. **Effect size as a key factor:** Replicability is primarily influenced by the true, unbiased effect size, highlighting the importance of targeting stable and reliable phenotypes.
4. **Promise of -based multivariate associations:** DWI-based brain-behaviour models should focus on phenotypes that display a sufficient temporal stability and test-retest reliability.

## MAIN

There is a growing interest in establishing and elucidating the intricate links between individual differences in phenotypic traits and measures of brain structure and function through brain-wide association studies (BWAS). Despite their potential, BWAS have been warranted to face challenges related to replicability and, in many cases, require large samples to be reproducible (Marek et al., 2022). In contrast to mass univariate BWAS, multivariate statistical learning techniques can achieve much higher effect sizes (Davatzikos, 2019; Woo et al., 2017) and can therefore be replicable with low to moderate sample sizes in many cases (Botvinik-Nezer & Wager, 2023; Spisak et al., 2023). This has recently been demonstrated with anatomical and functional MRI data such as cortical thickness (Kharabian Masouleh et al., 2019; Spisak et al., 2023), and functional connectivity (Spisak et al., 2023). However, a notable gap exists in our knowledge about the replicability of multivariate BWAS based on diffusion-weighted (DWI) MRI data. As DWI remains the primary non-invasive technique to offer insights into the microstructural integrity of neural pathways, replicability benchmarks for various DWI-based measures like fractional anisotropy (FA), axial (AD), mean (MD) and radial diffusivity (RD), as well as fibre tracking-based structural connectivity, are urgently needed to be able to draw informed decisions when planning, funding and reviewing future studies. Previous studies have reported that DWI-based multivariate models can deliver substantial predictive effect sizes, explaining up to 4-10% of variance in certain behavioural and psychometric phenotypes (Ooi et al., 2022), which – by theoretical considerations (supplementary information 1) – should lead to successful replicability with sample sizes in the hundreds. It is unclear, however, if such theoretical replicability estimates hold for real data (Macdonald, 2005).

Here we report the results of a systematic, large-scale, resampling-based empirical replicability analysis of DWI-based multivariate BWAS, including models based on streamline-based connectomes and structural properties such as FA, AD, MD, and RD, in case of 58 behavioural and psychometric phenotypes from the Human Connectome Project (Van Essen et al., 2013; Van Essen et al., 2012).

## METHODS IN BRIEF

We processed the high-quality multi-shell DWI data of N=901 participants available in the Human Connectome Project with FreeSurfer and MRtrix3 (see *supplementary information 2* and analysis code: *https://github.com/pni-lab/dwi-replicability*). We obtained a widely used set of cortical regions of interest with FreeSurfer via the Desikan-Killiany cerebral cortex atlas (Desikan et al., 2006) and generated the tractography between each pair of regions (supplementary information 2 for more details) with MRtrix3. We also obtained region-wise averages of five commonly used DWI-based measures, namely fractional anisotropy (FA), radial diffusivity (RD), axial diffusivity (AD) and apparent diffusion coefficient (ADC) and streamline-based structural connectivity (SC). These measures served as features in a multivariate machine learning analysis and aiming to predict 58 different behavioural phenotypes (supplementary information 3).

Replicability was estimated in line with previous work (Spisak et al., 2023). The data was randomly split into equally sized discovery and replication sets with 100 repetitions (n_rep_). Predictive models were trained on the discovery set separately for each phenotype with nested cross-validation. We used a simple but established modelling approach (Hoerl & Kennard, 1970), linear regression with L2 regularization; a.k.a Ridge, and employed a nested cross-validation approach for model development and evaluation in the discovery sample. The inner loop consisted of a 5-fold cross-validation optimizing the regularization parameter alpha (on a logarithmic space between 10^−4^ - 10^4^). Discovery effect size (explained variance) was obtained via the outer loop with another 5-fold cross-validation. The models (for each phenotype) were then finalized by training them on the whole discovery sample (without cross-validation). Next, we applied the finalized models on the replication set to predict phenotypes. This step did not entail model fitting; predictions were simply obtained as the dot product of replication-set features and the coefficients of the finalized models, as obtained during model discovery. We assessed the replicability of these predictions across all 100 discovery-replication splits as previously reported (Spisak et al., 2023), by calculating the probability of replicating a finding at a p-value threshold of 0.05 given that it was found to be significant in the discovery set (see equation on f*igure 1A*). The above computations were repeated for various sample sizes (25 – 450 samples in step size of 25) resulting in a total of ∼20 million model fits and thousands of core-hours of pre-processing and model trainings. Computations were performed on a high-performance computer cluster at the Institute for Artificial Intelligence in Medicine (IKIM), University Medicine Essen, Essen, Germany. We obtained analytical replicability estimates with Killeen’s formula (see *supplementary information 1 and 4*), based on effect sizes observed in the discovery phase and contrasted them with the empirical replicabilities.

## KEY FINDINGS

Our main findings are summarized in *figure 1* and *supplementary information 4*. In terms of predictive effect sizes, we corroborated previous results (Ooi et al., 2022) and found that effect sizes (in terms of R^2^) for replicable behavioural phenotypes ranged from 1.5% (cognitive fluid component) - 51.7% (for gender). We found that, as expected, the number of samples needed for replicability was inversely proportional to the logarithm of overall predictive effect sizes (measures in R^2^, r=-0.75). Empirical replicabilities did not deviate too much from theoretical effect size-based estimates. The ratio of replicable phenotypes (out of the 58 included phenotypes) at n≈450 was comparable across the investigated DWI measures (streamline-based connectivity, FA, RD, AD, ADC) and ranged between 24-31%, with streamline-based connectomes and FA providing the highest and lowest number of replicable phenotypes, respectively. Age and gender were highly predictable (100%) from all DWI-measures and replicable with sample sizes as low as 175 and 75, respectively. Among all DWI-based models, those predicting variables from the cognitive behavioural phenotype battery were found to be the most replicable (8 to 10 of 15 phenotypes with n<=450, 53-67%), followed by the motor battery (3/4, 75%). The emotion-related questionnaire battery was poorly replicable (1 to 2 of 22, 5-10%). None of the personality traits were found to be replicable with n<=450, neither were the two phenotypes from the alertness battery (MMSE, PSQI). Minimum number of samples required to reach the replicability criterion (P_replicability_>0.8) were lowest with streamline-based connectomes (median across all phenotypes that replicated with n<450: N≈163), followed by FA, RD, AD (all with N≈275), and ADC (N≈288).

**Figure 1.**
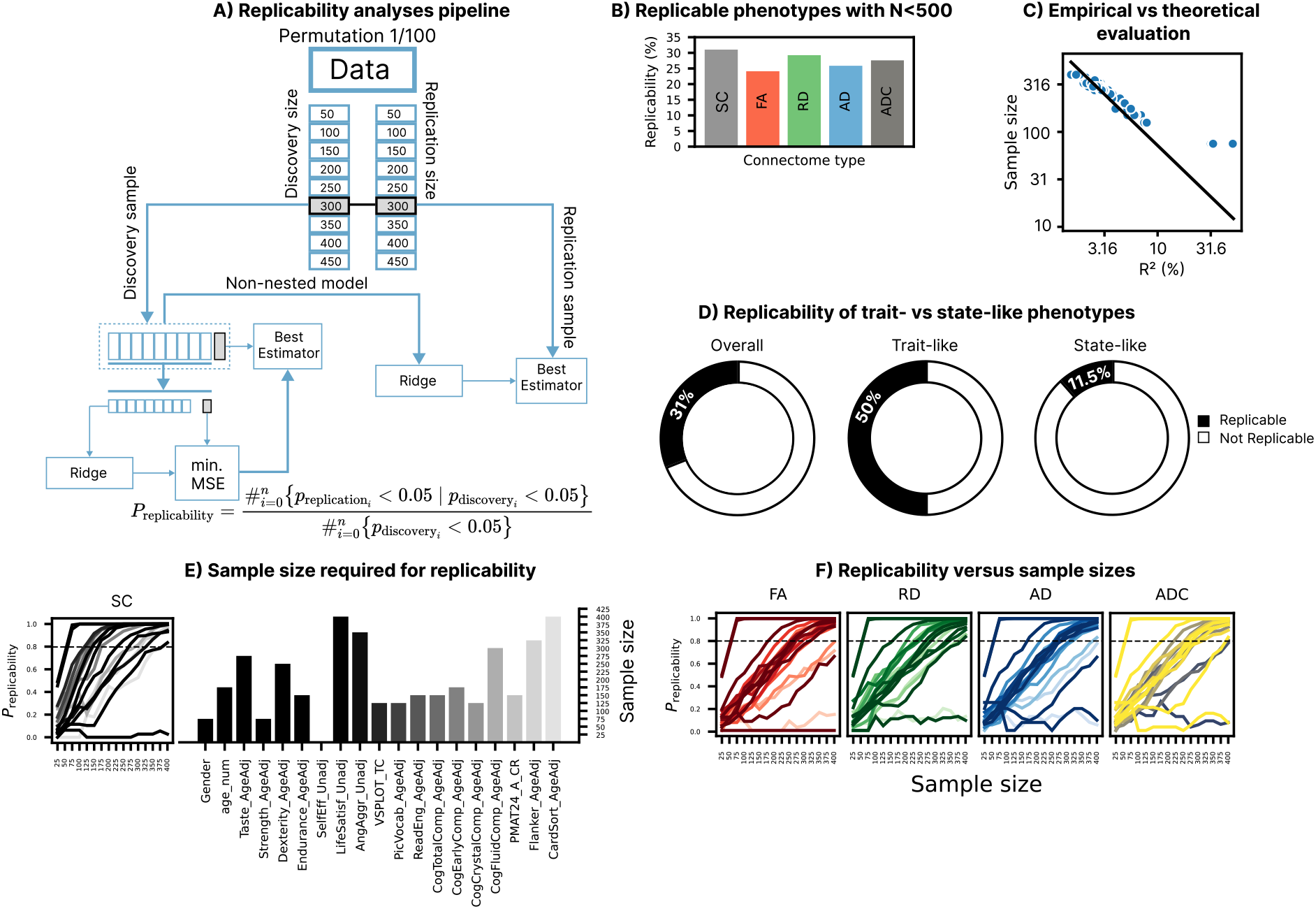
Replicability of DWI-predictive models of behaviour with small-to-medium sample sizes. **(A)** Machine learning pipeline for computing replicabilities of DWI-models. Data was permuted and split to equally sized discovery and replication sets. From both sets we randomly sampled variously sized samples. We trained a multivariate model in the discovery sample and evaluated its performance with nested cross-validation. The model was tested independently in the replication sample. Successful replication was defined as the probability of getting significant result in the replication sample, given that it was significant during discovery. The approach was repeated 100 times for each DWI-feature and phenotype. **(B)** Number of phenotypes with n<500 (out of 58 investigated phenotypes) with multivariate models based on streamline-based connectome, fractional anisotropy (FA), radial diffusivity (RD), axial diffusivity (AD) and apparent diffusion coefficient (ADC). Overall ∼30% of the included phenotypes were replicable with n<500. **(C)** The scatter plot shows the inverse relationship between effect sizes (R^2^) and the minimum sample size required for replicability (log_10_ scale). Points denote individual models while the line shows the theoretical estimates, calculated with Killeen’s replication probability (supplementary information 1). Empirical results and theoretical values are in good correspondence, except for phenotypes with high effect sizes (gender and strength). **(D)** Overall replicabilities across all connectome measures along with its trait- and state-like replicabilities. **(E-F)** Empirical replication probabilities of the investigated DWI-based brain features as a function of sample size (line plots) for different connectome-based models. The bar plot represents the minimum sample size required of achieving P_rep_>0.8 per phenotype.

## DISCUSSION

Our results, show that sample size requirements for replicable DWI-based BWAS are generally better than those observed with anatomical MRI data but worse than in case of resting state functional connectivity. For instance, to train a replicable model for the NIH total cognitive ability score n=75, 120 and 200 samples are needed with functional (resting state), diffusion-weighted and anatomical (cortical thickness) data, respectively (data for anatomical and functional data taken from (Spisak et al., 2023)). Similar to reports about functional and anatomical measures (Marek et al., 2022; Ooi et al., 2022; Spisak et al., 2023), we observed that a relatively high proportion of the investigated brain-phenotype models were either not significant or not replicable with the maximum investigated sample size (n=450). This sobering result warrants that the choice of the appropriate target phenotype is a crucial requirement for replicable BWAS. The question of ‘what is an appropriate target phenotype’ can be discussed from multiple aspects.

First, it is important to consider whether the phenotype of interest is relevant at all to the investigated brain measure. DWI-based brain features primarily capture microstructural characteristics of brain tissue, that are only subject to slow changes (due to development or plasticity) and are less prone to rapid alterations or fluctuations over time. Thus, we can expect that DWI data can only predict those measures reliably that are also persistent in time. From this perspective, phenotypes related to cognition, motor skills and demographics (e.g. age and gender) are well-suited to be predicted from DWI due to their inherent stability and/or trait-like nature. Indeed, we observed that large number of phenotypes belonging to such categories (labelled as ‘cognition’, ‘motor’, etc. in the HCP, see *supplementary information 3*) replicated with n<500.

In contrast, phenotypic measures related to emotions (accounts to ∼40% of all phenotypes in our study) and, to some degree even personality traits of openness and neuroticism, are known to be influenced by a variety of dynamic factors, from belief updates to contextual information or biological rhythms and cycles (Sharot et al., 2023; Thornton, Weaverdyck, Mildner, et al., 2019; Thornton, Weaverdyck, & Tamir, 2019), and may exhibit relatively rapid changes over time. DWI is inherently ill-posed to predict such phenotypes, explaining the observed poor replicabilities (5-10%).

Second, our results substantiate that – in line with theoretical considerations (see *supplementary information 1 and 4*) – the replicability of multivariate DWI-based brain-behaviour models are primarily a function of their effect sizes. This implies that, as substantiated by Gratton et al. (2022) and others (Ooi et al., 2024; Spisak et al., 2023), there are “two paths towards reliability”. Next to increasing sample sizes (possibly to the thousands), the reliability of the target phenotype, and its relevance to the brain measure, is also of central importance to improve the replicability. Phenotypes that are not only highly relevant to the brain feature of interest (e.g. trait measures for DWI), but also display a high test-retest reliability can be expected to display the highest effect sizes and will be replicable with small sample sizes. This has been exemplified by several previously reported multivariate neural signatures that showed strong, replicable effect sizes with various behavioural parameters, even though they were trained on small samples, sometimes fewer than 50 participants (Spisak et al., 2020; Wager et al., 2013; Woo et al., 2017). We argue that such models are the most promising candidates for individual-level predictions guiding clinical care and personalised medical approaches and our findings hint that the effect sizes and replicability necessary for these applications in not impossible to achieve, at least for the most reliable and stable phenotypes.

To achieve the necessary performance with DWI-based BWAS, the focus needs to be set on, (i) creating better phenotype measures and (ii) applying better modelling and feature processing methods. We believe that our results are conservative in this sense. On one hand, the investigated phenotypes were not chosen with DWI-based models in mind and, as discussed above, many of these phenotypes are simply not well suited for this purpose due to their state-like nature. On the other hand, we intentionally applied a minimalistic feature processing and modelling technique (Ridge regression with hyperparameter optimization), to keep results comparable to previous work and to avoid unnecessary analytic complexity for the presented, computationally already very intense, analyses. Better behavioural and psychometric phenotypes and advanced feature processing and modelling techniques (e.g. meta-matching, percolation analysis, ensemble learning) have the potential to improve reliability and effect sizes, but whether they do, remains to be seen.

Another significant challenge on the road towards replicable, and clinically useful, DWI-based models of individuals across datasets is between-site differences (Vollmar et al., 2010; Zhu et al., 2011) and dataset shift (Dockès et al., 2021). While our investigation is based on a homogenous dataset, careful MRI-sequence standardization measures and data harmonization approaches (Hu et al., 2023) promise to bridge this gap in the future and significantly improve out-of-centre generalizability of DWI-based brain-phenotype models.

## CONCLUSION

In conclusion, our study highlights that, with appropriate phenotype measures DWI-based multivariate BWAS can achieve moderate to high effect sizes and, consequently, deliver replicable associations with sample sizes in the hundreds. We argue that research that aims to provide clinically useful DWI-based brain-behaviour models should focus on phenotypes that display a sufficient temporal stability and test-retest reliability. Despite the promising results with such phenotypes, we must be aware of the substantial challenges ahead, including establishing broad generalizability across contexts, equity across subpopulations, and models with high neuroscientific validity and interpretability (Li et al., 2022; Wu et al., 2022). Addressing these challenges will require high precision target measures, large, representative samples and innovative new methods to process and model DWI data.

