## Supplementary material for "On the replicability of diffusion weighted MRI-based brain-behavior models"

***Supplementary information 1***

Analytical sample size calculations

Killeen’s replication probability (p_rep_) estimates the probability that a replication of an effect in a new sample will yield an effect of the same sign (no sign error). It depends on the effect size and degrees of freedom in the initial sample and the degrees of freedom in the replication sample and can be calculated from the observed t-value in the initial sample, or the P-value with an additional assumption of known variance. The probability of obtaining a significant result in a replication sample at a given statistical threshold (e.g., p < 0.05) is termed p_srep_, and is a straightforward extension. According to Eq. 6 from Lecoutre et al. (2010), Killeen’s probability of significant replication^6^ (p_srep_) can be extended to unknown variance:

$$p_{srep}(\alpha)=Pr \left( t_{rep}>T_{\alpha} \right| t_{obs})= Pr[{K^{'}}_{\left( v,v \right)}(t_{obs}/\sqrt{2})>T_{\alpha}/\sqrt{2}]$$

where K’ is the K-prime distribution with its parameter v set to the degrees of freedom, t_obs_ and t_rep_ are the observed student-t values in the discovery and replication samples, respectively and T_α_ is the t-value threshold that must be surpassed for a successful replication at an allowable false positive rate (significance-level) alpha. K-prime is a generalization of the noncentral t distribution, and in our case can be calculated from the initial sample t-value, degrees of freedom, and desired alpha level.

If statistical inference is based on Pearson’s correlation (r; as e.g. in Marek et al. (2022); Ooi et al. (2022)), the t-value for t_obs_ can be calculated as:

$$t_{obs}=r*\sqrt{\frac{n-2}{1-r^{2}}}$$

Based on the above equations, even if the correlation observed in the discovery sample is as low as r=0.1 (1% explained variance), at a sample size of n=1000, the probability of significant replication in a replication sample of the same size is:

$$p_{srep}=0.86$$

From the same equation, sample sizes to achieve 80% replication probability with r=0.19 and r=0.32 and (4 to 10% variance explained, as reported for DWI measures in Ooi et al. (2022)) are n=269 and 91, respectively.

***Supplementary information 2***

Magnetic Resonance Imaging protocol

Detailed information regarding the T1-weighted and diffusion weighted sequences are mentioned elsewhere (https://www.humanconnectome.org/, HCP_S1200_Release_Reference_Manual.pdf). In brief, T1-weighted scans are acquired with repetition time 2400 milli seconds (ms), echo time 2.14 ms, inversion time 1000 ms, flip angle 8^0^, field of view 224 × 224 mm, voxel size 0.7 mm isotropic, bandwidth 210 H_Z_/P_X_, iPAT (integrated Parallel Acquisition Techniques) 2, and acquisition time 7 minutes 40 seconds. The diffusion weighted images are acquired in 90 directions, consisting of an equal number of shells i.e., b=1000, b=2000, and b=3000 s/mm^2^, along with 6 b=0 images. The diffusion weighted parameters included repetition time 5520 ms, echo time 89.5 ms, flip angle 78^0^, refocusing flip angle 160^0^, field of view 210 × 180 mm, matrix 168 × 144, slice thickness 1.25 mm, 111 slices, 1.25 mm isotropic voxel, multiband factor 3, echo spacing 0.78 ms, bandwidth 1488 H_Z_/P_X_, phase partial Fourier 6/8, right-to-left and left-to-right phase encoding polarities.

Image processing

Image data analyses were conducted for achieving participant level structural connectivity matrices – connectomes – which are comprised of nodes and edges. Nodes were defined as gray matter regions of interest, and edges connect a pair of nodes through white matter tractography. For the nodes, individual anatomical T1-weighted images were segmented into bilateral regions of comprising of 68 nodes based on Desikan-Killiany-Tourville atlas (Klein & Tourville, 2012) using FreeSurfer. Diffusion weighted image analyses were conducted using MRtrix3 to generate whole brain white matter-based tractography. Tractography analyses were performed incorporating the recommendations provided in MRtrix3 basic and advanced tractography tutorial (https://osf.io/fkyht/). In brief, images from both phase encoding directions (right-to-left and left-to-right) were denoised (*dwidenoise*) and corrected for Gibbs ringing artefacts (*mrdegibbs*). Subsequently, corrections for motion, eddy-current and susceptibility induced distortions were applied using the *dwifslpreproc* command, which invokes FSL’s eddy and topup processes. Response functions for white matter, gray matter, and cerebrospinal fluid were estimated using the dwi2response dhollander command. Fiber Orientation Distributions (FOD) within each voxel were estimated with multi-shell multi-tissue constrained spherical deconvolution (*dwi2fod msmt_csd*), utilizing the diffusion data and response functions. The FODs were then normalized using mtnormalise. The T1 image was segmented into five distinct tissue classes (cortical and subcortical gray matter, white matter, cerebrospinal fluid, and pathological tissue) using *5ttgen* and co-registered to the diffusion space with the b=0 image serving as the reference. Subsequently, an interface image of the gray-white matter boundary was generated using *5tt2gmwmi*. 10 million streamlines were generated with *tckgen*, based on the white matter FOD image, guided by the anatomic constraints of the 5tt image, and seeded from the GM-WM interface. These tractograms were filtered using the *tcksift2* algorithm, which normalizes the tractogram by assigning a cross-sectional area multiplier, aiming to obtain biologically accurate measures of fiber connectivity. Finally, using *tck2connectome*, DWI-based connectomes were generated for connectivity strength (scaling with the inverse of node volumes). Along with the connectivity strength, average matrices for diffusion tensor maps namely fractional anisotropy (FA), radial diffusivity (RD), apparent diffusion coefficient (ADC) and axial diffusivity (AD) were also calculated, which reflects the microstructural properties of white matter connection integrities.

Replicability assessment machine learning pipeline

For the replicability analyses (see *figure 1A and* [*https://github.com/pni-lab/dwi-replicability*](https://github.com/pni-lab/dwi-replicability)), connectomes were considered as features, with behavioral measures which served as different targets. Machine learning models were constructed with linear regression (Ridge/L2 regularization). Data was equally split into two samples, discovery sample (for train-test) and replication sample (validation). The models were developed on the discovery sample using a nested cross-validation approach with 5-fold split at both outer and inner loops of the cross validation. Subsequently, the models were applied to both the discovery and replication sets for generating predictions of phenotypes, and the p-values of both discovery and replication datasets were determined (i.e., p-value for y_discovery and yhat_discovery, y_replication and yhat_replication). This model pipeline was repeated with 100 shuffles, for each phenotype and replication probability per phenotype was calculated at p-value <0.05. In addition, the model results were also obtained for variable sample sizes, i.e., 25-450 in step size of 25. For a given sample size, and number of permutations, n=100, replication probability was calculated as:

P _replicability_ = $\frac{{\#}_{i=0}^{n} \left\{ {p_{replication}}_{i}<0.05 | {p_{discovery}}_{i}<0.05 \right\}}{{\#}_{i=0}^{n}\{p_{{discovery}_{i}}<0.05\}}$

Where octothorp #{} denotes the number of permutation cases satisfying condition within the curly brackets and, p_discovery_ is the p-value for the cross-validated model performance estimate during discovery, p_replication_ is the p-value of the association between model output and target phenotype in the replication sample and P_replicability_ is the probability for a given behavioural model to be replicable at a given sample size.

***Supplementary information 3***

Phenotypes – domains and subdomains

| Domain | Subdomain | Score [serial number] |
| --- | --- | --- |
| Alertness | Cognitive Status (Mini Mental Status Exam) | MMSE_Score [1] |
|  | Sleep (Pittsburgh Sleep Quality Index) | PSQI_Score [2] |
| Cognition | Episodic Memory (Picture Sequence Memory) | PicSeq_AgeAdj [3], AgeAdj = age adjusted |
|  | Executive Function/Cognitive Flexibility (Dimensional Change Card Sort) | CardSort_AgeAdj [4] |
|  | Executive Function/Inhibition (Flanker Inhibitory Control and Attention Task) | Flanker_AgeAdj [5] |
|  | Fluid Intelligence (Penn Progressive Matrices) | PMAT24_A_CR [6], CogFluidComp_AgeAdj [7], CogCrystalComp_AgeAdj [8], CogEarlyComp_AgeAdj [9], CogTotalComp_AgeAdj [10] |
|  | Language/Reading Decoding (Oral Reading Recognition) | ReadEng_AgeAdj [11] |
|  | Language/Vocabulary Comprehension (Picture Vocabulary) | PicVocab_AgeAdj [12] |
|  | Processing Speed (Pattern Comparison Processing Speed) | ProcSpeed_AgeAdj [13] |
|  | Self-regulation/Impulsivity (Delay Discounting) | DDisc_AUC_200 [14] |
|  | Spatial Orientation (Variable Short Penn Line Orientation Test) | VSPLOT_TC [15] |
|  | Sustained Attention (Short Penn Continuous Performance Test) | SCPT_SEN [16], SCPT_SPEC [17] |
|  | Memory\|Verbal Episodic Memory (Penn Word Memory Test) | IWRD_TOT [18] |
|  | Working Memory (List Sorting) | ListSort_AgeAdj [19] |
| Emotion | Emotion Recognition (Penn Emotion Recognition Test) | ER40HAP (happiness)[20], ER40NOE (neutral) [21], ER40SAD (sadness) [22], ER40FEAR (fear) [23], ER40ANG (anger) [24] |
|  | Negative Affect (Sadness, Fear, Anger) | AngAffect_Unadj [25], AngHostil_Unadj [26], AngAggr_Unadj [27], FearAffect_Unadj [28], FearSomat_Unadj [29], Sadness_Unadj [30] |
|  | Psychological Well-being (Positive Affect, Life Satisfaction, Meaning and Purpose) | LifeSatisf_Unadj [31], MeanPurp_Unadj [32], PosAffect_Unadj [33] |
|  | Social Relationships and Social Developments | Friendship_Unadj [34], Loneliness_Unadj [35], PercHostil_Unadj [36], PercReject_Unadj [37], EmotSupp_Unadj [38], InstruSupp_Unadj [39] |
|  | Stress and Self-Efficacy (Perceived Stress, Self-Efficacy) | PercStress_Unadj [40], SelfEff_Unadj [41] |
| Motor | Endurance (2-minute walk test) | Endurance_AgeAdj [42] |
|  | Locomotion (4-meter walk test) | GaitSpeed_Comp [43] |
|  | Dexterity (9-hole PegBoard) | Dexterity_AgeAdj [44] |
|  | Strength (Grip Strength Dynamometry) | Strength_AgeAdj [45] |
| Personality | Five Factor Model (NEO-FFI) | NEOFAC_A [46], NEOFAC_O [47],  NEOFAC_C [48], NEOFAC_N [49],  NEOFAC_E [50] |
| Sensory | Audition (Words in Noise) | Noise_Comp [51] |
|  | Olfaction (Odor Identification Test) | Odor_AgeAdj [52] |
|  | Pain (Pain Intenstiy and Interference Surveys) | PainIntens_RawScore [53],  PainInterf_Tscore [54] |
|  | Taste (Regional Taste Intensity Test) | Taste_AgeAdj [55] |
|  | Contrast Sensitivity (Mars Contrast Sensitivity) | Mars_Final [56] |
| Demography | Age and Gender | Age [57], Gender [58] |

***Supplementary table 4***

Empirical effect sizes given a replicability value to achieve (> 80%).

|  | Effect sizes as measured in R^2^ | | | | | Empirical sample sizes for achieving > 80% | | | | | Theoretical sample sizes for achieving > 80% | | | | |
| --- | --- | --- | --- | --- | --- | --- | --- | --- | --- | --- | --- | --- | --- | --- | --- |
| **Phenotypes** | **Connectivity** | **FA** | **RD** | **AD** | **ADC** | **Connectivity** | **FA** | **RD** | **AD** | **ADC** | **Connectivity** | **FA** | **RD** | **AD** | **ADC** |
| **CardSort_AgeAdj** | 1.710761 | - | - | - | - | 400 | - | - | - | - | 466 | - | - | - | - |
| **Flanker_AgeAdj** | 2.577560 | - | 2.105100 | - | - | 325 | - | 375 | - | - | 308 | - | 378 | - | - |
| **PMAT24_A_CR** | 6.966490 | 3.019982 | 2.927795 | 2.794531 | 2.948202 | 150 | 300 | 325 | 325 | 300 | 111 | 262 | 270 | 284 | 269 |
| **CogFluidComp_AgeAdj** | 2.456813 | - | 1.953462 | 1.531179 | 1.736997 | 300 | - | 375 | 400 | 400 | 323 | - | 407 | 521 | 459 |
| **CogCrystalComp_AgeAdj** | 8.013802 | 3.328193 | 3.762440 | 3.091602 | 3.431250 | 125 | 275 | 250 | 275 | 275 | 96 | 237 | 210 | 256 | 230 |
| **CogEarlyComp_AgeAdj** | 4.012146 | 2.127961 | 2.572881 | 2.008739 | 2.231151 | 175 | 325 | 275 | 325 | 300 | 196 | 374 | 308 | 396 | 356 |
| **CogTotalComp_AgeAdj** | 6.249368 | 3.720116 | 4.427224 | 3.655241 | 3.729355 | 150 | 225 | 225 | 225 | 225 | 125 | 212 | 177 | 216 | 211 |
| **ReadEng_AgeAdj** | 5.190035 | 2.640197 | 2.634399 | 2.526705 | 2.640322 | 150 | 300 | 275 | 325 | 300 | 151 | 300 | 301 | 314 | 300 |
| **PicVocab_AgeAdj** | 7.473113 | 2.922465 | 3.413821 | 2.693796 | 3.093113 | 125 | 275 | 250 | 275 | 275 | 103 | 271 | 231 | 294 | 256 |
| **VSPLOT_TC** | 7.902811 | 3.031311 | 3.497828 | 3.007412 | 3.086421 | 125 | 275 | 250 | 275 | 275 | 98 | 261 | 226 | 263 | 256 |
| **AngAggr_Unadj** | 2.452190 | 2.914258 | 2.082865 | 2.985861 | 2.140405 | 350 | 275 | 350 | 275 | 325 | 324 | 272 | 382 | 265 | 371 |
| **LifeSatisf_Unadj** | 1.897191 | - | - | - | - | 400 | - | - | - | - | 420 | - | - | - | - |
| **SelfEff_Unadj** | - | - | 1.844334 | - | 1.710466 | - | - | 400 | - | 400 | - | - | 432 | - | 466 |
| **Endurance_AgeAdj** | 6.044950 | 4.831483 | 5.343023 | 5.177830 | 4.904573 | 150 | 200 | 175 | 175 | 200 | 129 | 162 | 146 | 151 | 160 |
| **Dexterity_AgeAdj** | 3.597771 | 3.151380 | 2.757204 | 2.867365 | 2.797003 | 250 | 275 | 300 | 300 | 300 | 219 | 251 | 287 | 276 | 283 |
| **Strength_AgeAdj** | 31.429247 | 31.639403 | 32.827863 | 33.569361 | 33.271692 | 75 | 75 | 75 | 75 | 75 | 21 | 21 | 20 | 20 | 20 |
| **Taste_AgeAdj** | 2.552265 | 2.883204 | 3.101147 | 2.895066 | 2.974010 | 275 | 275 | 275 | 275 | 275 | 311 | 275 | 255 | 274 | 266 |
| **Age** | 5.606442 | 2.382677 | 2.563374 | 2.681055 | 2.473375 | 175 | 325 | 350 | 300 | 300 | 139 | 333 | 309 | 296 | 321 |
| **Gender** | 51.714579 | 49.780590 | 50.465796 | 51.038111 | 50.969116 | 75 | 75 | 75 | 75 | 75 | 11 | 12 | 12 | 12 | 12 |

***References***

Klein, A., & Tourville, J. (2012). 101 labeled brain images and a consistent human cortical labeling protocol. *Frontiers in neuroscience*, *6*, 33392.

Lecoutre, B., Lecoutre, M.-P., & Poitevineau, J. (2010). Killeen's probability of replication and predictive probabilities: How to compute, use, and interpret them. *Psychological Methods*, *15*(2), 158.

Marek, S., Tervo-Clemmens, B., Calabro, F. J., Montez, D. F., Kay, B. P., Hatoum, A. S., Donohue, M. R., Foran, W., Miller, R. L., & Hendrickson, T. J. (2022). Reproducible brain-wide association studies require thousands of individuals. *Nature*, *603*(7902), 654-660.

Ooi, L. Q. R., Chen, J., Zhang, S., Kong, R., Tam, A., Li, J., Dhamala, E., Zhou, J. H., Holmes, A. J., & Yeo, B. T. (2022). Comparison of individualized behavioral predictions across anatomical, diffusion and functional connectivity MRI. *Neuroimage*, *263*, 119636.
